# Modified Synchronization Modulation Electric Field attenuates acute brain injury and improves neurological outcomes following stroke induced by transient middle cerebral artery occlusion in rats

**DOI:** 10.64898/2026.11.21.768295

**Authors:** David Raj Chellappan, Nadezhda N. Zheleznova, Zhi Tian, Nathan Hall, Kristof Williams, Georgina Agyekum, Yuqian Tian, Wei Chen, Lianchun Wang, Kanita Beba Abadal, Ruisheng Liu

**Affiliations:** Department of Molecular Pharmacology and Physiology, Morsani College of Medicine, University of South Florida, Tampa, FL 33612, USA; WR Biotech, LLC, Tampa, FL, 33612, USA; Department of Neurology, University of South Florida Morsani College of Medicine, Tampa, FL, USA; USF Health Transplant Research Center (UTRC), University of South Florida, Tampa, FL, 33612, USA

**Keywords:** Cerebral ischemia–reperfusion, middle cerebral artery occlusion, synchronization modulation electric field, Na⁺/K⁺-ATPase, neuroprotection

## Abstract

Cerebral ischemia disrupts energy-dependent ion transport and causes acute brain injury. The synchronization modulation electric field (SMEF) platform was developed to support Na⁺/K⁺-ATPase function under conditions of limited ATP availability and has shown protective effects in renal ischemia models. We investigated whether cerebral cortical application of third-generation SMEF attenuates cerebral ischemia–reperfusion injury in rats. Adult male Sprague–Dawley rats underwent sham surgery or transient middle cerebral artery occlusion (MCAO) for 120 min, with or without SMEF treatment during ischemia. At 24 h after reperfusion, body-weight measurements, neurological deficit, open-field locomotor activity, 2,3,5-triphenyltetrazolium chloride (TTC) based cerebral infarct volume, hemispheric swelling, cortical and striatal histopathology, and Western blot analysis of Na⁺/K⁺-ATPase α1, glial fibrillary acidic protein (GFAP) and nuclear factor-kappa B (NF-κB). Compared with untreated MCAO rats, SMEF-treated rats exhibited significantly less body-weight loss, lower neurological deficits, greater total distance of movement, smaller edema-corrected infarct volumes and percentages, and lower cortical and striatal injury scores. SMEF also shifted Na⁺/K⁺-ATPase α1, GFAP and NF-κB protein abundance toward control levels. These findings support an acute protective effect of SMEF following transient MCAO. Further studies should assess Na⁺/K⁺-ATPase activity and longer-term functional, histological and molecular outcomes.

## Introduction

Stroke is a leading cause of mortality and long-term neurological disability worldwide. According to the Global Burden of Disease Study 2021, approximately 11.9 million new strokes occurred worldwide in 2021, and 93.8 million people were living with a history of stroke. Stroke accounted for approximately 7.3 million deaths and 160.5 million disability-adjusted life-years in 2021. Ischemic stroke was the most common subtype globally in 2021, accounting for 65.3 % of incident strokes globally, with the highest proportion observed in high-income countries (Feigin et al., 2025; GBD 2021 Stroke Risk Factor Collaborators, 2024). Ischemic stroke occurs when cerebral blood flow is interrupted or severely reduced, depriving brain tissue of oxygen and metabolic substrates. Restoration of blood flow is essential for salvaging viable tissue; however, reperfusion can also exacerbate cellular and molecular injury. Cerebral ischemia-reperfusion injury involves interrelated processes, including energy failure, ionic imbalance, excitotoxicity, oxidative stress, mitochondrial dysfunction, neuroinflammation, blood–brain barrier (BBB) disruption and cerebral edema, which collectively contribute to neuronal death (Qin et al., 2022). The neurovascular unit comprises neurons, glial cells, brain microvascular endothelial cells, pericytes and extracellular matrix components, which collectively maintain BBB integrity and cerebral homeostasis (Gong et al., 2025). Damage to this unit, including loss of BBB integrity, facilitates the infiltration of circulating inflammatory cells and extravasation of plasma-derived components into the ischemic brain, further aggravating tissue injury (Levinson et al., 2025).

Na⁺/K⁺-ATPase is an ATP-dependent membrane transporter that maintains the transmembrane sodium and potassium gradients essential for maintenance of the resting membrane potential, neuronal excitability, cell volume regulation, and secondary active transport (Clausen et al., 2017). During cerebral ischemia, ATP depletion compromises Na⁺/K⁺-ATPase function and disrupts ionic homeostasis, contributing to membrane depolarization, cellular swelling, calcium dysregulation and excitotoxic injury (Khanna et al., 2014; Song and Yu, 2014). Huang et al. (2015) reported reduced Na⁺/K⁺-ATPase activity in the ischemic penumbra following transient middle cerebral artery occlusion (MCAO) in rats. Furthermore, Li et al. (2021) implicated the interaction between Na⁺/K⁺-ATPase and glutamate transporter-1 (GLT-1) in protection against cerebral ischemia–reperfusion injury. Together, these findings provide a rationale for investigating strategies aimed at preserving Na⁺/K⁺-ATPase function to mitigate ischemic brain injury.

Cerebral ischemia also induces astrocytic reactivity, which is commonly associated with changes in glial fibrillary acidic protein (GFAP) expression. Reactive astrocytes can influence neuroinflammatory signaling, BBB integrity, cerebral edema, while also contributing to tissue repair and glial scar formation at later stages (Patabendige et al., 2021; Shen et al., 2021). In addition to astrocytic reactivity, nuclear factor kappa B (NF-κB) signaling contributes to the inflammatory response following cerebral ischemia-reperfusion. Experimental studies have associated increased NF-κB p65 expression with elevated levels of pro-inflammatory cytokines, including tumor necrosis factor – α and interleukin-6, in the ischemic brain (Li et al., 2023). Assessment of GFAP and NF-κB expression therefore provides complementary molecular endpoints for examining whether SMEF treatment is associated with changes in inflammatory signaling and astrocytic reactivity following cerebral ischemia-reperfusion.

Our laboratory developed the synchronization modulation electric field (SMEF) platform to synchronize and modulate Na⁺/K⁺-ATPase function. In previous studies, SMEF sustained Na⁺/K⁺-ATPase currents in ATP-depleted muscle fibers and maintained the transepithelial potential difference in isolated kidneys. SMEF also delayed ATP depletion and preserved mitochondrial respiratory function during renal ischemia. *In vivo*, SMEF reduced injury in mouse models of renal ischemia-reperfusion and kidney transplantation (Chen et al., 2022). Recently, a preprint from our laboratory reported that the modified third-generation SMEF (3m-SMEF) reduced hypoxia/reoxygenation-induced cell death in HK-2 cells and attenuated renal ischemia-reperfusion injury in mice. These protective were accompanied by preservation of renal ATP and Na⁺/K⁺-ATPase expression and localization (Hall et al., 2026). These findings further support investigating SMEF-based approaches for protection against ischemic injury in other organs, including the brain. Collectively, these findings provide a mechanistic rationale for extending SMEF approach to cerebral ischemia, in which ATP depletion and impaired Na⁺/K⁺-ATPase function contributes to the loss of ionic gradients, membrane depolarization and cellular injury. We therefore hypothesized that the cortical application of modified SMEF during transient MCAO would attenuate ischemic brain injury and improve neurological outcomes after reperfusion.

In the present study, we investigated the neuroprotective effects of SMEF treatment in a rat model of transient MCAO. Therapeutic efficacy was evaluated using complementary structural, functional, histopathological, and molecular endpoints, including TTC-based infarct volume and cerebral swelling measurements, neurological deficit assessment, open-field locomotor activity, hematoxylin and eosin staining, and Western blot analysis of Na⁺/K⁺-ATPase α1, GFAP and NF-κB. We aimed to determine whether SMEF treatment attenuates acute ischemic brain injury and functional impairment after transient focal cerebral ischemia and to explore whether these effects are associated with preservation of Na⁺/K⁺-ATPase α1 expression and changes in GFAP and NF-κB expression.

## Materials and methods Animals and ethical approval

Adult male Sprague-Dawley (SD) rats (8 -10 weeks old; 250-300 g) were obtained from Envigo and acclimatized to the animal facility for at least 7 days before experimentation. SD rats were housed in groups of two to three per cage under controlled environmental conditions (temperature, 22 ± 2⁰C; relative humidity, 30-70%; 12-h light/12-h dark cycle) with *ad libitum* access to standard laboratory chow and water throughout the study. Animals were randomly assigned to the experimental groups, and all the behavioral assessments, histological evaluation and biochemical analyses were performed by investigators blinded to treatment allocation. All the experimental procedures were performed under protocol 11894R, approved by the University of South Florida Institutional Animal Care and Use Committee (IACUC), and followed National Institutes of Health (NIH) guidelines for laboratory animal care and use.

## Experimental design and study groups

Rats were randomly assigned to three experimental groups: sham, MCAO and MCAO + SMEF (Figure 1). Sham-operated rats underwent electrode implantation and expose cervical blood vessels without MCA occlusion or SMEF stimulation. Rats in the MCAO and MCAO + SMEF groups underwent 120 min of MCAO followed by 24 h reperfusion. In the MCAO + SMEF group, stimulation was applied throughout the ischemic period and discontinued at the onset of reperfusion. Body weight, neurological deficits, and spontaneous locomotor activity were assessed at 24 h after reperfusion. Brain tissues were collected for TTC-based infarct measurement, histopathological evaluation and Western blot analysis.

**Figure 1.**
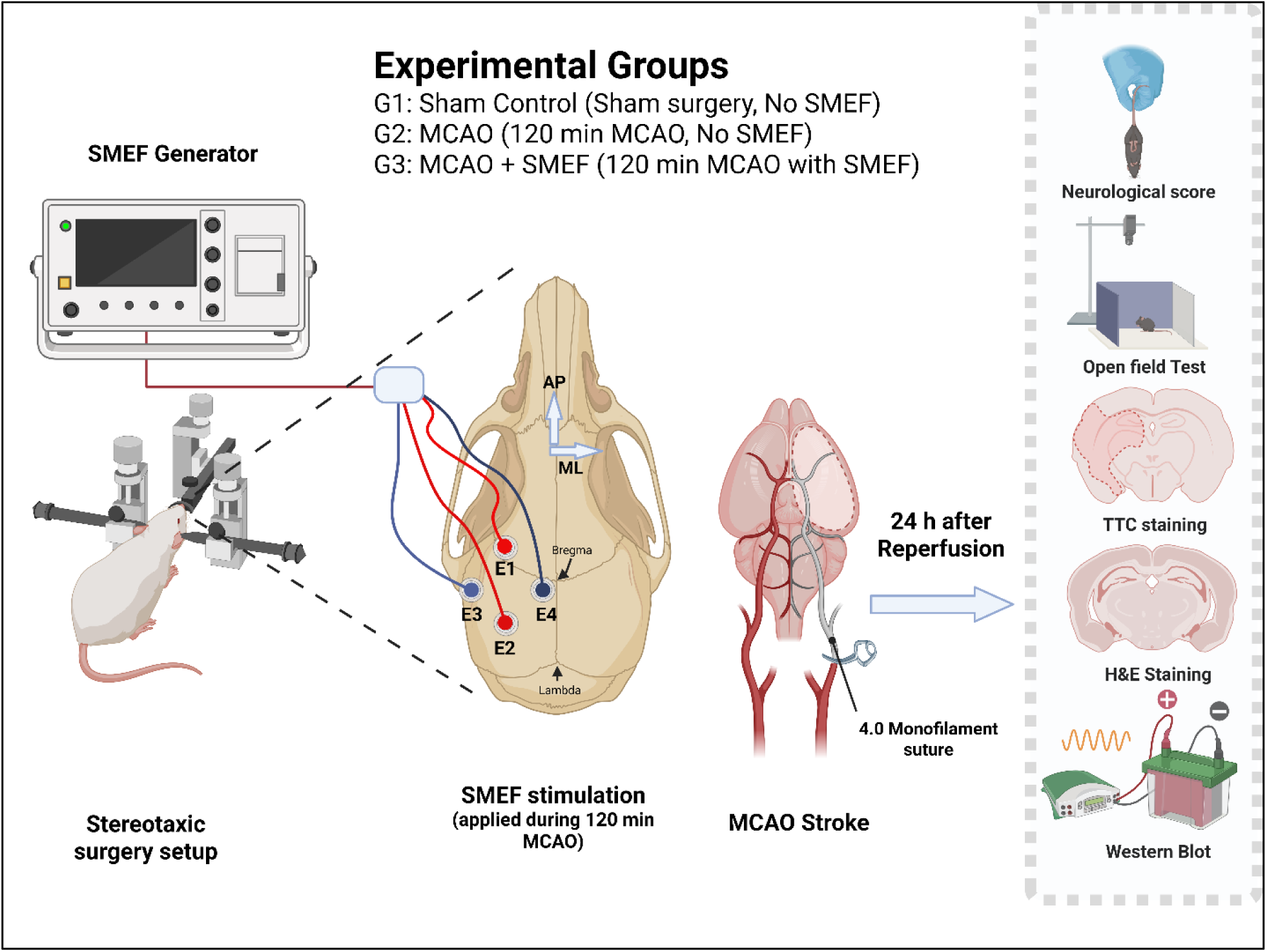
Experimental design and SMEF treatment in a rat model of transient MCAO. Rats were assigned to sham, MCAO, and MCAO + SMEF groups. Four electrodes (E1-E4) were implanted in the left cerebral hemisphere, ipsilateral to MCA occlusion to deliver SMEF through two orthogonal electrode pairs. In the treatment group, SMEF was applied during the 120-min MCAO period and discontinued at the onset of reperfusion. Neurological function and open-field locomotor activity were assessed at 24 h after reperfusion, and brain tissues were collected for TTC staining, histopathology, and Western blot analysis. Created in BioRender. Chellappan, D. R. (2026) https://BioRender.com/w8agw7a

## Stereotaxic electrode implantation

SD rats were anesthetized with 3-4% isoflurane for induction and maintained under anesthesia with 1.5 – 2.0% isoflurane delivered in an oxygen-containing carrier gas. Throughout surgery, body temperature was monitored continuously with a rectal probe and maintained at 37.0 ± 0.5°C with a feedback-controlled heating system (ATC-2000; WPI). Rats were weighed before surgery and at the study endpoint. Ophthalmic ointment was applied to both eyes to prevent corneal drying during anesthesia, and lidocaine solution was injected at the incision sites to provide local analgesia.

Each rat was positioned in a stereotaxic frame (RWD Instruments) equipped with an anesthesia mask, and the head was secured with ear bar and an incisor bar. The scalp hair was trimmed, and the surgical site was disinfected with 10% povidone-iodine solution equivalent to 1% available iodine (Dynarex corporation, NY, USA). Bupivacaine (0.2 ml of a 0.25% solution) was injected subcutaneously along the scalp midline before incision to provide local anesthesia during electrode implantation (Vöröslakos et al., 2021). A 1.5-2 cm midline incision was made to expose the dorsal surface of the skull, and the periosteum was gently removed using a cotton swab. After the skull was cleaned and dried, bregma and lambda were identified. Bregma was used as the reference point for anteroposterior (AP) and mediolateral (ML) coordinates (Paxinos et al., 1985; Schwarz et al., 2010).

Four electrodes were implanted in the left cerebral hemisphere, ipsilateral to the MCA occlusion, at the following coordinates relative to bregma: E1, AP +2.34 mm and ML -2.31 mm; E2, AP −2.34 mm and ML -2.31 mm; E3, AP −0.74 mm and ML -4.31 mm; and E4, AP −0.74 mm and ML -0.35 mm. Negative ML values indicate positions left to the midline. A burr hole was drilled at each marked site using a microdrill with sterile saline irrigation to minimize the mechanical and thermal injury to the underlying dura and cortical tissues.

Each electrode consisted of a female socket (E363/0; Protech International, USA) crimped onto a stainless-steel wire of diameter 127 µm (A-M systems, WA, USA). The socket remained outside the skull, with its supporting edge resting on the skull surface and the wire extending 2 mm below this edge. The electrodes were inserted through the burr holes into the cortex, with cortical penetration estimated at approximately 1.0-1.2 mm. The same electrode configuration was used across all animals to standardize insertion length relative to the skull surface.

The electrodes were secured to the skull using acrylic dental cement prepared by mixing fast-curing acrylic resin liquid with auto-polymerizing acrylic resin powder (Jet Liquid and Jet XR powder). The dental cement was allowed to harden completely, the implantation sites were inspected for bleeding or other complications, and the scalp incision was closed around the electrode assembly using 4-0 monofilament nylon sutures (662G, Ethicon, Somerville, NJ, USA) and the triple antibiotic ointment (Trifecta Pharmaceuticals, FL, USA) was applied to the incision area (Schwarz et al., 2010). The electrode assembly was then connected to the synchronization modulation electric field (SMEF) generator using male plug contacts (E363/A; Protech International, USA).

Following electrode implantation, each rat was positioned supine, and the ventral neck region was shaved, disinfected and prepared aseptically for MCAO surgery. Sham-operated rats underwent the same electrode implantation procedure but did not receive SMEF stimulation.

## Transient Middle Cerebral Artery Occlusion (MCAO) stroke

Transient focal cerebral ischemia was established by occluding the left middle cerebral artery (MCA) with an intraluminal monofilament, using a modification describe by Longa et al. (1989). The cervical region was accessed through a midline incision, after which the submandibular glands were gently separated and retracted. The left common carotid artery (CCA), external carotid artery (ECA), and internal carotid artery (ICA) were then exposed and carefully dissected free from adjacent connective tissue and the vagus nerve. The pterygopalatine artery (PPA) was temporarily ligated close to its origin using an 8-0 nylon suture during filament placement and ischemia. The distal ECA was identified, cauterized and divided with microsurgical scissors. The CCA and ICA were temporarily occluded with vascular clips.

An arteriotomy was made in the ECA stump, and a 4-0 silicone-coated nylon monofilament, with total length 4 cm, silicon-coated head-end diameter 0.38-0.40 mm, suture body diameter 5-6 mm, coating length 0.20 mm (MSRC40B200PK50, RWD Life science, China) was introduced into the ICA and advanced toward the origin of the middle cerebral artery (MCA). The filament was gently advanced until resistance was encountered, consistent with placement at the MCA origin. The monofilament was secured at the ECA stump using an 8-0 suture and left in place for 120 min to induce transient focal cerebral ischemia.

In the SMEF-treated group, stimulation was applied throughout the 120-min ischemic period. During ischemia, body temperature was monitored continuously with a rectal probe and maintained at 37.0 ± 0.5°C using a feedback-controlled heating system. After 120 minutes, the SMEF generator was turned off, and the filament was carefully withdrawn to initiate reperfusion. The ligature around the pterygopalatine artery was then removed. The ECA stump was secured and inspected for bleeding, and the muscular layer was sutured using 4-0 braided polyglactin sutures (VCP496G, Ethicon, Somerville, NJ, USA). Finally, the midline cervical incision was closed with 4-0 monofilament nylon sutures (662G, Ethicon, Somerville, NJ, USA) and the triple-antibiotic ointment (Trifecta Pharmaceuticals, FL, USA) was applied to the incision site.

After surgery, rats recovered in a temperature-controlled recovery cage and were monitored for return of consciousness and mobility, neurological status, wound condition, and access to food and water. Meloxicam (5 mg/kg per dose) was administered subcutaneously before surgery and again after surgery for perioperative analgesia. Softened food and water gel were placed within easy reach. All rats received warmed, sterile 0.9% normal saline (1 mL/100 g body weight) subcutaneously to support postoperative hydration (S. Liu et al., 2009; Oh and Narver, 2024).

Sham-operated rats underwent a midline cervical incision and isolation of the corresponding blood vessels without vascular occlusion. All these rats received the same analgesic regimen and saline supplementation as the other experimental groups.

## Modified SMEF stimulation protocol

To improve the efficacy of the third-generation synchronization modulation electric field platform, we modified the field application strategy, waveform, and stimulation parameters. The modified platform incorporates dual energy transformation functions and power injections designed to enhance ATP generation. The pumping rate and synchronization of pump molecules were further optimized to minimize potential adverse cellular effects. Because neuronal and glial cell membranes have diverse orientations within the brain, the interaction between the applied electric field and membrane-associated Na⁺/K⁺-ATPase may vary with membrane orientation. To improve coverage of the targeted cortical region, four electrodes arranged as two orthogonal pairs were used to apply electric fields sequentially along the anteroposterior and mediolateral axes, alternating every three stimulation cycles. This approach was intended to increase the probability of synchronizing Na⁺/K⁺-ATPase molecules within the stimulated tissue.

The SMEF protocol comprised three phases: synchronization, modulation and maintenance. The synchronization phase was designed to synchronize the cycling of Na^+^/K^+^-ATPase molecules with the applied electrical field. The activation pulse was set to 300 mV for 200 µs, the energy-trap pulse to 300 mV for 400 µs, and the plateau amplitude to 150 mV. The initial and target frequencies were set to 40 and 100 Hz, respectively, with a frequency increment of 2%. The synchronization pulse count was set to 300. During the modulation phase, the stimulation frequency was gradually adjusted to the predetermined targeted frequency. This frequency was then maintained for the remaining 120-min cerebral ischemic period.

This synchronization-and-acceleration strategy was designed to support restoration of transmembrane ionic gradients and membrane potential during ischemia while reducing dependence on limited intracellular ATP, thereby promoting preservation of cerebral cellular function. The SMEF generator was configured for continuous operation without a preset time limit, and SMEF was applied during the 120-min cerebral ischemic period and discontinued before reperfusion. Anesthesia was maintained with 1.5-2% isoflurane during stimulation.

## Modified Bederson neurological score

Neurological function was assessed 24h after reperfusion using a modified Bederson neurological deficit scoring system (Bederson et al., 1986c; Crumrine et al., 2012). All assessments were performed by an investigator blinded to the treatment allocation. Each rat was placed individually on the flat surface and allowed to move freely to assess the spontaneous circling and walking ability. Forelimb flexion was evaluated by gently lifting the rat by the base of the tail and observing the forelimb extension. Resistance to lateral push was assessed by applying gentle lateral pressure and observing the rat’s ability to maintain its posture. Neurological deficits were scored on a 0-4 scale as follows; 0, no observable neurological deficit and normal walking; 1, contralateral forelimb flexion or failure to fully extend the contralateral forelimb; 2, forelimb flexion accompanied by decreased resistance to lateral push; 3, decreased resistance to lateral push accompanied by spontaneous circling towards the paretic side and; 4, inability to walk spontaneously. Higher scores indicated greater neurological impairment.

## Open field test

Spontaneous locomotor activity was evaluated using the open-field test at 24 h after reperfusion following transient MCAO. Each rat was placed individually in the center of an open-field arena (75 x 50 x 35 cm) and allowed to explore freely for 4 min. Testing was conducted in a quiet environment under consistent conditions across animals. Animal movements were recorded throughout the session using a digital camera mounted 100 cm above the arena to capture its entire interior of the open field. The arena was cleaned between trials to minimize residual olfactory cues. Animal movements were video-recorded throughout the session and subsequently analyzed using ANY-maze video-tracking software (version 7.65), with identical arena boundaries defined for all trials. The total distance traveled during the 4-min session was calculated for each rat as a measure of spontaneous locomotor activity (Li et al., 2013).

## TTC staining and infarct volume analysis

After 24 h of reperfusion, rats were deeply anaesthetized and transcardially perfused with ice-cold physiological saline to remove the residual blood. Following perfusion, brains were rapidly excised, placed on a chilled surface, and cut into six consecutive 2-mm thick coronal slices using a prechilled rat brain matrix. The brain slices were incubated in 2% 2,3,5-triphenyltetrazolium chloride solution (TTC; T8877; Sigma-Aldrich) at 37 °C for approximately 20 minutes in the dark, as previously described with modifications (Bederson et al., 1986a; Ryang et al., 2011).

Following TTC staining, the slices were rinsed and fixed in 4% paraformaldehyde (PFA) for 24 h before imaging. The anterior and posterior surfaces of each slice were digitally scanned using a color image scanner (CanoScan LiDE 400, Canon, USA). Viable tissue stained red, whereas the infarcted tissue appeared pale or unstained. Images were spatially calibrated and analyzed using ImageJ (NIH, Bethesda, MD, USA) by an investigator blinded to group allocation. On each surface, the infarcted area and the total ipsilateral and contralateral hemisphere areas were manually delineated. The non-infarcted ipsilateral area was calculated by subtracting the directly measured infarcted area from the total ipsilateral area.

For each slice, the anterior and posterior measurements of each area were averaged and multiplied by the slice thickness (2 mm) to obtain the corresponding volumes. These volumes were summed across all six slices for each animal. Edema-corrected infarct volume was calculated using the indirect method by subtracting the non-infarcted ipsilateral hemisphere volume from the contralateral hemisphere volume. The corrected infarct volume was expressed as a percentage of the contralateral hemisphere volume. (S. Liu et al., 2009; Swanson et al., 1990). Hemisphere swelling (%) was calculated by subtracting the contralateral hemisphere volume from the ipsilateral hemisphere volume, dividing the difference by the contralateral hemisphere volume, and multiplying by 100 (Wang et al., 2015).

## Histopathological assessment

At 24 h after reperfusion, brain tissue samples collected were fixed in 4 % PFA for 24 h and subsequently immersed in 70% ethanol for an additional 24 h. paraffin embedding and sectioning were performed at the H. Lee Moffitt Cancer Center and Research Institute. Coronal sections (4 µm) thick were stained with hematoxylin and eosin (H&E) using a standard protocol. Whole section images were acquired using an Olympus VS120 slide scanning system (Olympus, Tokyo, Japan) at the Lisa Muma Weitz Imaging Core Facility, University of South Florida and examined at magnifications ranging from 4x to 40x.

Histological injury was assessed using a study-specific semiquantitative scoring system comprising neuronal injury (0-4) and tissue vacuolation/rarefaction (0-3). The component scores were summed to obtain a composite injury score ranging from 0 to 7, with higher scores indicating greater injury severity. Detailed scoring criteria are provided in supplementary Table S1.

## Western Blot Analysis

After 24h of MCAO, brain tissue collected from the ipsilateral cerebral region was homogenized on ice in RIPA lysis buffer (catalog no. 9806; Cell Signaling Technology) supplemented with protease inhibitor cocktail (catalog no. 87785; Thermo Scientific). Tissue homogenates were incubated on ice with intermittent mixing and subsequently centrifuged at 12,000 × g for 15 min at 4°C. The supernatant was collected, and total protein concentration was determined using a bicinchoninic acid (BCA) protein assay kit (catalog no. 23227).

Equal amounts of protein (30 µg per lane) were mixed with Laemmli sample buffer containing a reducing agent and heated at 95°C for approximately 5 min. Proteins were separated by sodium dodecyl sulfate-polyacrylamide gel electrophoresis (SDS-PAGE) and transferred onto polyvinylidene difluoride (PVDF) membranes. Following transfer, membranes were blocked with 5% non-fat dry milk in Tris-buffered saline containing 0.1% Tween-20 (TBST) for 1 h at room temperature and then incubated overnight at 4°C with primary antibodies against GFAP (catalog no. 3670, Cell Signaling Technology), Na⁺/K⁺-ATPase α1 (catalog no. 58953T, Cell Signaling Technology), NF-kB p65 (catalog no. 16502, Abcam), β-Tublin (catalog no. AC030, ABclonal), and GAPDH (catalog no. 2118, Cell Signaling Technology). The approximate molecular weights of GFAP, Na⁺/K⁺-ATPase α1, NF-kB p65, β-Tubulin and GAPDH were 50, 100–110, 65, 55 and 37 kDa respectively.

Following primary antibody incubation, membranes were washed three times with TBST and incubated with horseradish peroxidase (HRP)-conjugated goat anti-rabbit IgG (1:10,000; A6154, Sigma-Aldrich) or goat anti-mouse IgG (1:5,000; A4416. Sigma-Aldrich), as appropriate, for 1 h at room temperature. After additional TBST washes, immunoreactive bands were visualized using an enhanced chemiluminescence (ECL) substrate (catalog no. 170-5060, Bio-Rad) and captured using a digital chemiluminescence imaging system (Universal Hood II, Bio-Rad Laboratories.

Band intensities were quantified by densitometric analysis using ImageJ 1.54p (NIH, USA). Background signal was subtracted consistently across all samples, and the intensity of each target protein was normalized to the corresponding GAPDH or β-Tubulin band intensity from the same sample. Normalized values were then divided by the mean normalized value of the sham group to express protein abundance as fold change relative to sham.

## Statistical analysis

Statistical analyses were performed using GraphPad Prism (version 10.5.0; GraphPad Software). Each animal was considered an independent experimental unit. Continuous data are presented as mean ± standard deviation (SD), whereas ordinal neurological deficit and histopathological injury scores are presented as median with interquartile range (IQR). Individual animal values are shown in the figures.

For continuous outcomes, normality was assessed using graphical inspection and the Shapiro–Wilk test. Comparisons between two independent groups were performed using an unpaired, two-tailed Student’s *t*-test when normality and equal-variance assumptions were reasonable, or Welch’s ANOVA followed by Dunnett’s T3 multiple-comparisons test was used. Continuous outcomes unsuitable for parametric analysis were compared using the Mann-Whitney U test for two groups or the Kruskal-Wallis test followed by Dunn’s multiple-comparisons test.

Ordinal neurological deficit scores were analyzed using the Kruskal–Wallis test followed by Dunn’s multiple-comparisons test. Histopathological injury scores were compared between MCAO and MCAO + SMEF groups using the Mann-Whitney *U* test. Percentage body-weight change was calculated for each animal as [(weight at 24 h after reperfusion − presurgical body weight)/presurgical body weight] × 100. Between group comparison of body-weight change were performed using one-way ANOVA followed by Tukey’s multiple-comparison test. All applicable tests were two-tailed. Multiplicity-adjusted *P* values were used for post hoc comparisons, and *P* < 0.05 was considered statistically significant. Sample sizes and the statistical tests used are reported in the corresponding figure legends.

## Results

### SMEF attenuated body-weight loss following transient MCAO

At 24 h after reperfusion, rats subjected to MCAO exhibited significantly greater body-weight loss than sham-operated rats (*P* < 0.0001; Fig. 2). SMEF treatment significantly attenuated this weight loss compared with the MCAO group (*P* < 0.0001). However, body-weight loss remained significantly greater in the MCAO + SMEF group than in the sham group (*P* < 0.001), indicating that SMEF partially attenuated acute postischemic body-weight loss.

**Figure 2.**
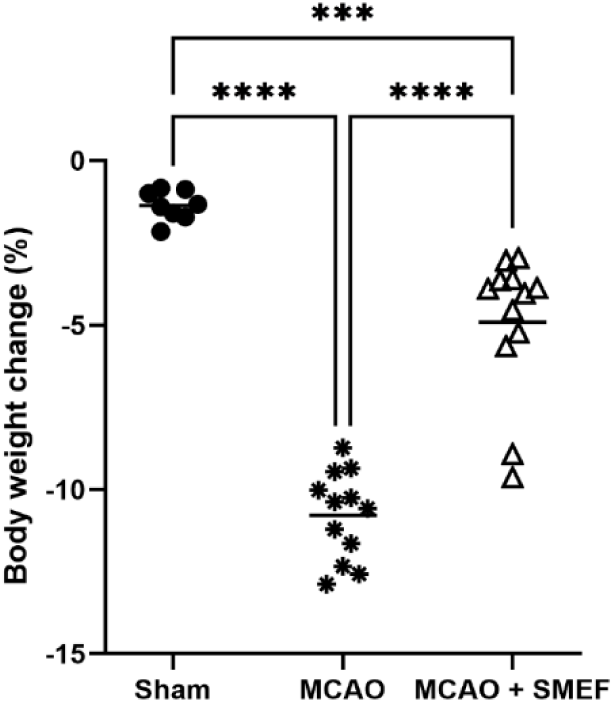
SMEF attenuates body-weight loss following transient MCAO. Body weight was measured before surgery and 24 h after reperfusion. Percentage body-weight change was calculated as [(body weight at 24 h after reperfusion – preoperative body weight)/preoperative body weight] × 100. Negative values indicate body-weight loss. Each symbol represents an individual animal. Data are presented as mean ± SD (n = 12 rats per group). Differences among groups were analyzed using one-way ANOVA, followed by Tukey’s multiple-comparisons test. ***P < 0.001 and ****P < 0.0001 for the comparisons indicated by brackets.

### SMEF reduced neurological deficits following transient MCAO

Neurological function was assessed using the modified Bederson neurological deficit scoring system at 24 h after reperfusion. Sham-operated rats exhibited no observable neurological deficits, whereas rats in the MCAO group had significantly higher neurological deficit scores, ranging from 3 to 4 (*P* < 0.001 vs. sham; Fig. 3). SMEF treatment significantly reduced neurological deficit scores compared with the untreated MCAO rats (*P* < 0.001), with neurological scores ranging from 2 to 3, indicating partial improvement in neurological function.

**Figure 3.**
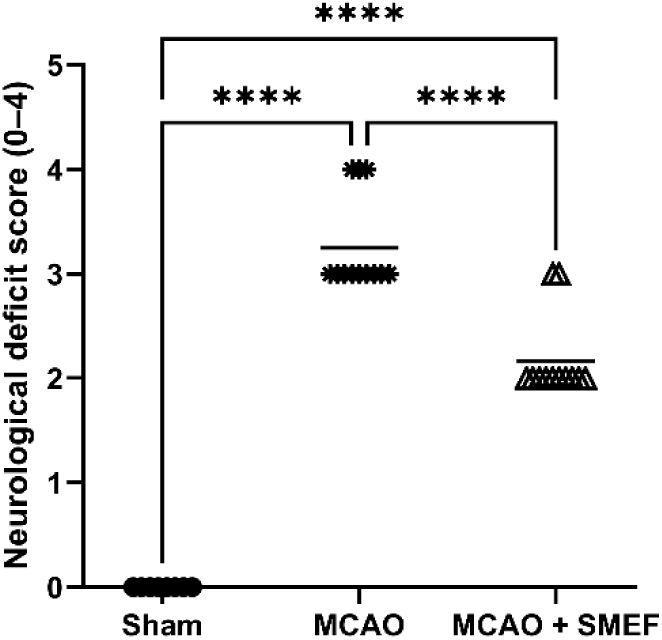
SMEF attenuates neurological deficits following transient MCAO. Neurological function was assessed 24 h after reperfusion using a modified Bederson neurological deficit scoring system (0–4), with higher scores indicating greater neurological impairment. Each symbol represents an individual animal, and the data are presented as mean ± SD (n = 12 rats per group). Difference among the sham, MCAO, and MCAO + SMEF groups were analyzed using the Kruskal-Wallis test followed by Dunn’s multiple-comparisons test. ****P < 0.0001 for the comparison indicated by brackets.

### SMEF reduced cerebral infarction following transient MCAO in rats

TTC staining at 24 h after reperfusion revealed extensive pale, unstained regions in the ipsilateral hemisphere of MCAO rats, whereas SMEF-treated rats exhibited smaller infarcted regions (Fig. 4A). Quantitative analysis showed that SMEF treatment significantly reduced both edema-corrected infarct volume and edema-corrected infarct percentage compared with MCAO group (both *P* < 0.01; Fig. 4B–C). Hemispheric swelling was numerically lower in the MCAO + SMEF group than in the MCAO group, although this difference was not statistically significant (*P* > 0.05; Fig. 4D).

**Figure 4.**
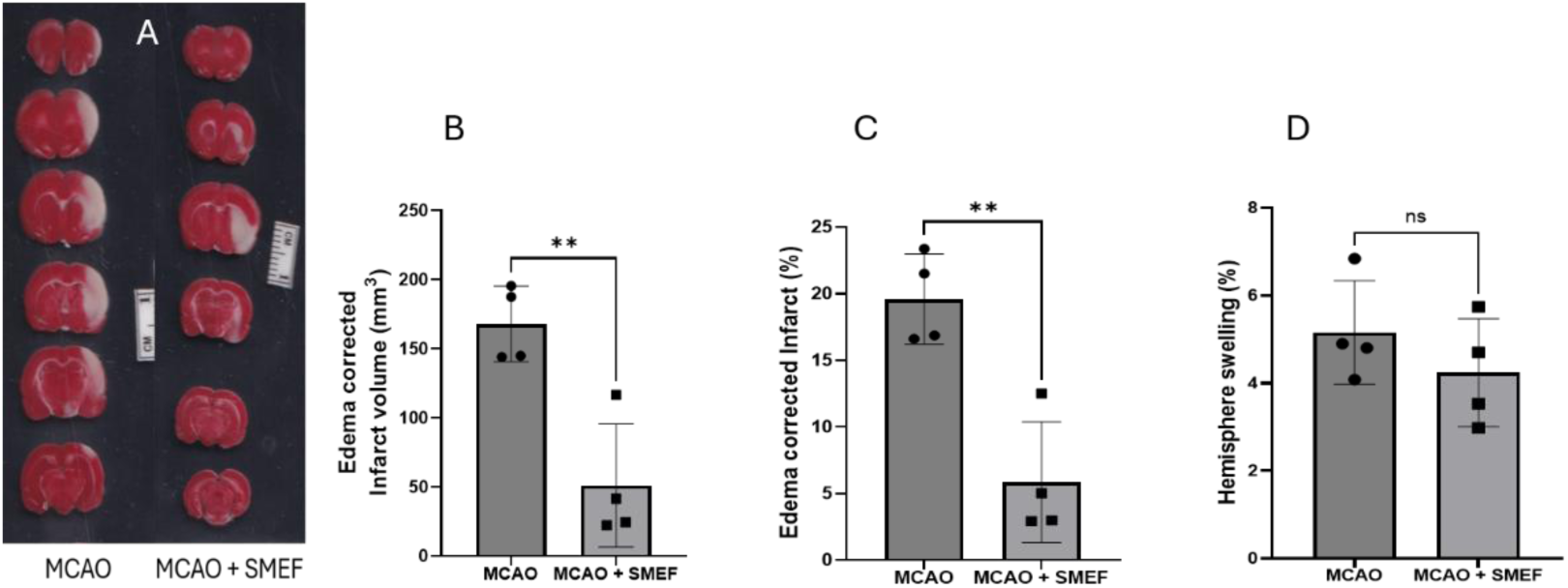
SMEF reduces cerebral infarction following transient middle cerebral artery occlusion (MCAO) in rats. (A) Representative TTC-stained coronal brain sections obtained 24 h after reperfusion from MCAO and MCAO + SMEF-treated rats. Viable brain tissue stained red, whereas infarcted tissue remained pale or unstained. Measurements from the anterior and posterior surfaces of each 2-mm-thick coronal section were averaged for volumetric analysis. (B) Edema-corrected infarct volume (mm³), calculated as the contralateral hemisphere volume minus the non-infarcted ipsilateral hemisphere volume. (C) Edema-corrected infarct volume expressed as a percentage of the contralateral hemisphere volume. (D) Hemispheric swelling (%), calculated from the difference between ipsilateral and contralateral hemisphere volumes relative to the contralateral hemisphere. Individual symbols represent individual animals, and data are presented as mean ± SD (n = 4 rats per group). Comparisons between MCAO and MCAO + SMEF groups were performed using an unpaired two-tailed Student’s t-test. P < 0.05 was considered statistically significant. **P < 0.001; ns, not statistically significant, for the comparisons indicated.

### SMEF improved spontaneous locomotor activity following transient MCAO

Spontaneous locomotor activity was assessed using the open-field test at 24 h after reperfusion. The total distance traveled during a 4-min session was quantified using automated video tracking. Rats in the MCAO + SMEF group traveled a significantly greater total distance than untreated MCAO rats in open-field at 24 h after reperfusion (*P* < 0.01; Fig. 5B), indicating improved spontaneous locomotor activity. Individual distances varied more widely in the MCAO + SMEF group than in the MCAO group.

**Figure 5.**
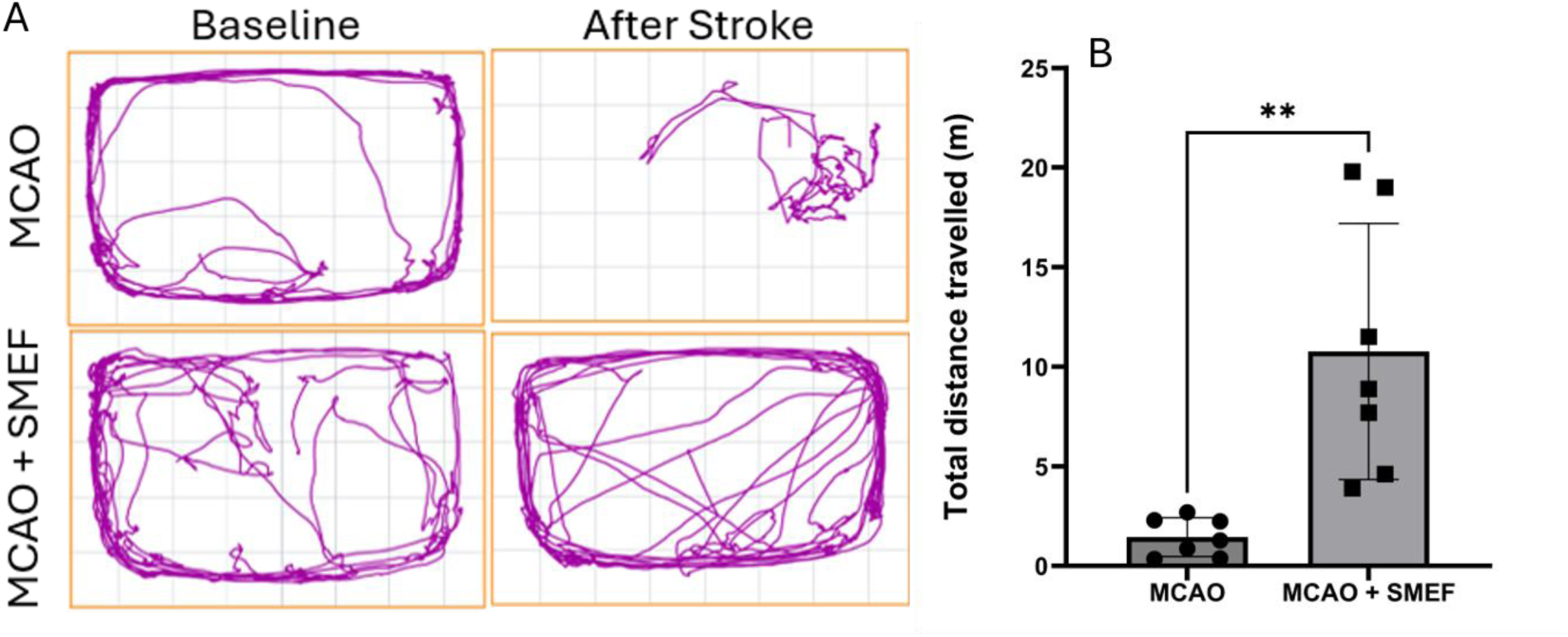
SMEF improves spontaneous locomotor activity following transient MCAO. (A) Representative open-field movement trajectories of MCAO and MCAO + SMEF-treated rats during a 4-min testing session at 24 h after reperfusion. (B) Total distance traveled during the 4-min session. Each symbol represents an individual animal. Data are presented as mean ± SD (n = 7 rats per group). Differences between groups were analyzed using an unpaired, two-tailed *t*-test. \*\**P* < 0.01 versus MCAO.

### SMEF reduced cortical and striatal histopathological injury scores following transient MCAO

At 24 h after reperfusion, untreated MCAO rats exhibited histological changes in the ipsilateral cortex and striatum, eosinophilic neurons characterized shrunken neurons with pyknotic nuclei and intensely eosinophilic cytoplasm, accompanied by neuropil vacuolation. These abnormalities were less pronounced in SMEF-treated rats, which showed comparatively preserved neuronal morphology and reduced vacuolation (Fig. 6A and 6B).

**Figure 6.**
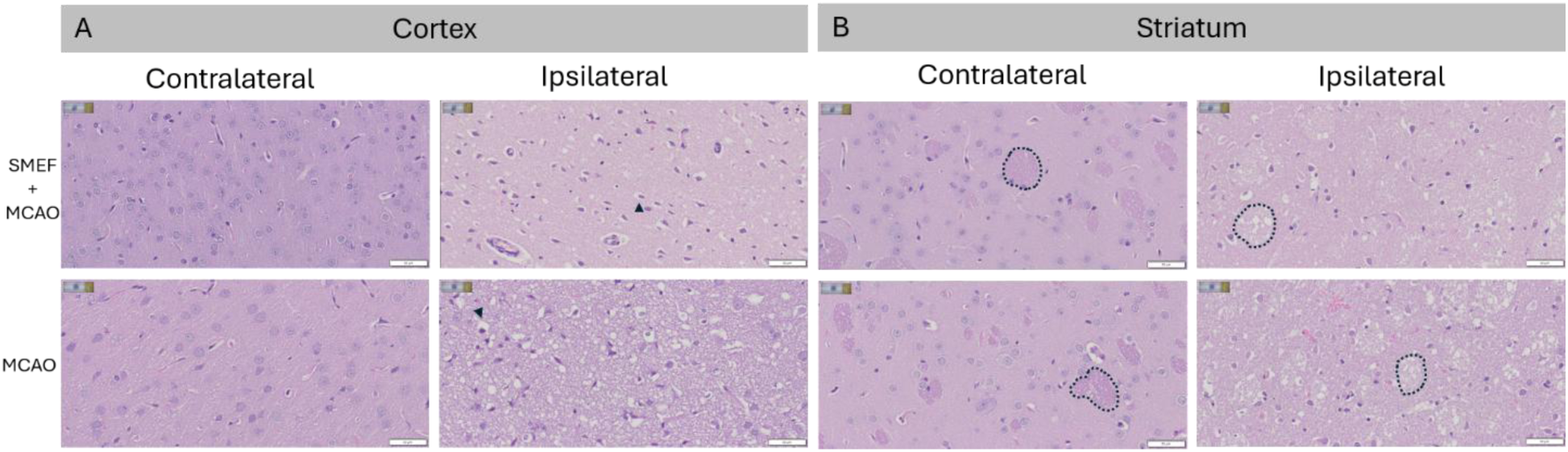
SMEF attenuates cortical and striatal histopathological injury following transient MCAO. Representative hematoxylin and eosin (H&E)-stained sections of the cortex (A) and striatum (B) from the ipsilateral and contralateral hemispheres of MCAO and MCAO + SMEF-treated rats at 24 h after reperfusion. The black arrowhead indicates a shrunken cell with a condensed, darkly stained nucleus, consistent with pyknosis. The dotted outline indicates striatal fiber bundles, with more prominent intra- bundle vacuolation in the ipsilateral than in the contralateral region of striatum. Scale bar, 50 μm.

Consistent with these observations, rats in the MCAO + SMEF group exhibited lower semiquantitative histopathological injury scores than untreated MCAO rats in both the cortex (Fig. 7A) and striatum (Fig. 7B). Residual histopathological abnormalities remained evident in both regions after SMEF treatment, indicating that tissue injury was attenuated but not completely prevented.

**Figure 7.**
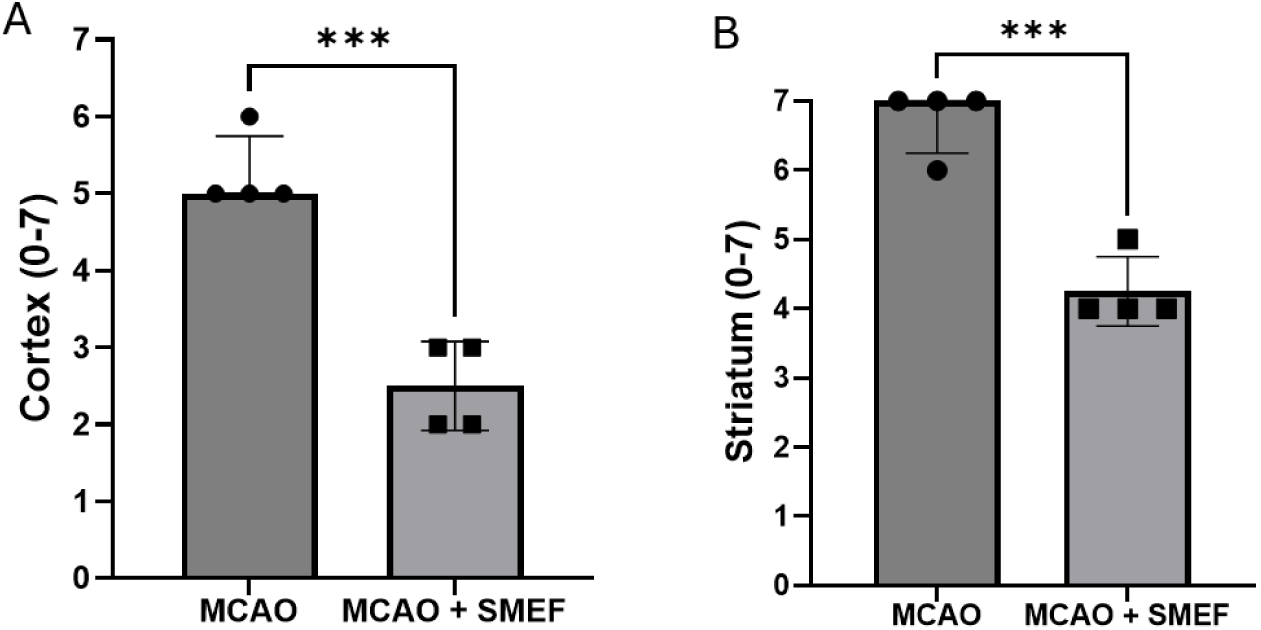
SMEF reduces cortical and striatal histopathological injury scores following transient MCAO. Semiquantitative histopathological injury scores in the cortex (A) and striatum (B) at 24 h after reperfusion. Composite scores ranged from 0 to 7 and comprised neuronal injury (0-4) and edema/vacuolation (0-3), with higher scores indicating greater tissue injury. Scoring criteria are provided in Supplementary Table S1. Each symbol represents an individual animal. Data are presented as median with interquartile range (IQR; n = 4 rats per group). Comparisons between MCAO and MCAO + SMEF groups were performed using the Mann–Whitney *U* test. *P* < 0.05 was considered statistically significant.

### SMEF modified Na^+^/K^+^-ATPase α1, GFAP and NF-κB p65 protein levels following transient MCAO

At 24 h after reperfusion, Western blot analysis revealed significantly higher GAPDH-normalized Na^+^/K^+^- ATPase α1protein levels in the MCAO stroke rats than in the sham group (*P* < 0.01; Fig. 8D). SMEF treatment significantly reduced Na^+^/K^+^-ATPase α1 levels compared with the untreated MCAO group (*P* < 0.05), with no statistically significant difference between the sham and MCAO + SMEF groups. In contrast, GAPDH-normalized GFAP protein levels were significantly lower in the MCAO group than in the sham group (*P* < 0.01; Fig. 8C). SMEF treatment significantly increased GFAP protein levels compared to the untreated MCAO group (*P* < 0.05), with no statistically significant difference between the sham and MCAO + SMEF groups.

**Figure 8.**
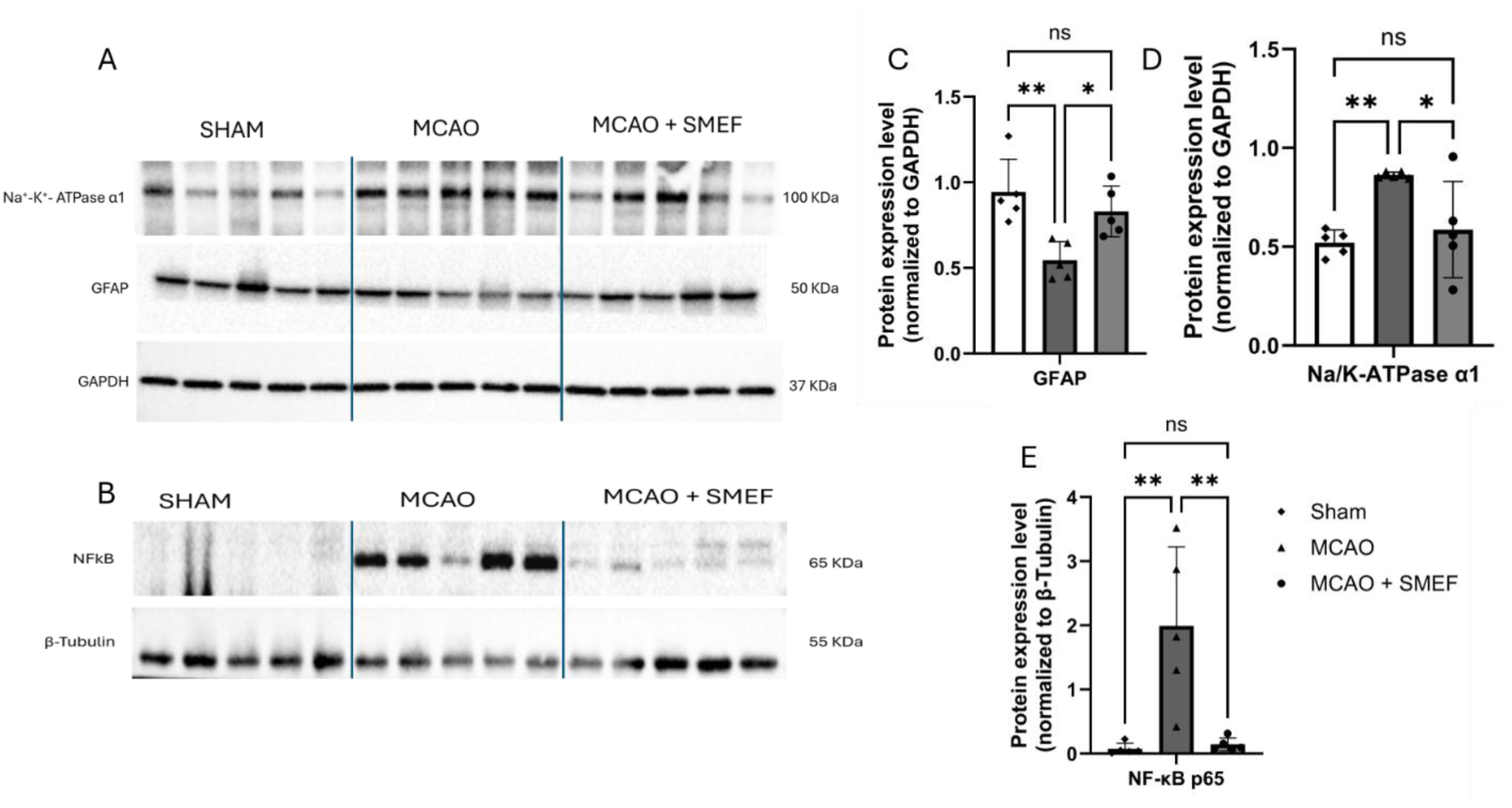
Effects of SMEF treatment on Na⁺/K⁺-ATPase α1, GFAP and NF-κB p65 protein levels following transient MCAO. Brain tissue was collected from sham, MCAO and MCAO + SMEF groups at 24 h after reperfusion. (A) Representative Western blots of Na⁺/K⁺-ATPase α1, GFAP, NF-κB p65, and GAPDH. (B) Representative Western blots of NF-κB p65 and β-tubulin. (C, D) Densitometric quantification of GFAP (C) and Na⁺/K⁺-ATPase α1 (D), normalized to GAPDH. (E) Densiometric quantification of NF-κB p65 normalized to β-tubulin. Each symbol represents an individual animal. Data are presented as mean ± SD (n = 5 rats per group). Group differences were analyzed using one-way ANOVA followed by Tukey’s multiple-comparison test. \**P* < 0.05; \*\**P* < 0.01; ns, not statistically significant, for the comparisons indicated by brackets.

Furthermore, β-tubulin-normalized NF-κB p65 protein levels were significantly higher in the MCAO group than in the sham group (*P* < 0.01; Fig. 8E). SMEF treatment significantly reduced NF-κB p65 levels compared with the untreated MCAO group (*P* < 0.01), with no statistically significant difference between the sham and MCAO + SMEF groups.

## Discussion

The present study extends our laboratory’s previous work on SMEF-mediated protection against renal ischemia–reperfusion injury to cerebral ischemia. The SMEF platform was developed to synchronize and modulate Na⁺/K⁺-ATPase function under conditions of limited ATP availability. In our previous study, third- generation SMEF sustained pump currents in ATP-depleted muscle fibers, maintained the transepithelial potential difference in isolated kidneys, and reduced renal injury in mouse ischemia–reperfusion and kidney transplantation models. SMEF also delayed ATP depletion and preserved mitochondrial respiratory function during renal ischemia (Chen et al., 2022). Collectively, these findings provided the rationale for investigating whether SMEF could attenuate cerebral ischemia-reperfusion injury. In the present study, cortical application of SMEF during the 120 min ischemic period reduced infarct burden and histopathological injury and improved neurological scores and spontaneous locomotor activity at 24 h after reperfusion. These findings support an acute protective effect in this rat model.

SMEF treatment during the ischemic phase attenuated postoperative body-weight loss compared with untreated MCAO rats, although weight loss remained greater than in sham-operated animals. This finding suggests a partial improvement in overall postoperative conditions. However, body weight loss is a non- specific outcome that may reflect reduced food and water intake, fluid imbalance and surgical stress. In filament MCAO procedures, ischemia of the external carotid artery territory can impair mastication and swallowing, restrict ingestion, and contribute to postoperative weight loss (Dittmar et al., 2003). The reduction in body weight at 24 h after reperfusion is consistent with the previous reports in the rats subjected to MCAO (Liu et al., 2011; Modo et al., 2000). They also reported a marked reduction in body weight at 24 h after MCAO. In addition, neurological deficits may also interfere with feeding and drinking. Consequently, changes in body weight cannot be attributed exclusively to cerebral injury. Because food consumption, water intake, and hydration status were not measured, the mechanisms underlying the attenuation of weight loss remain uncertain.

Neurological assessment and open-field testing provided complementary measures of acute functional outcome. Neurological examination complements measurements of infarction and tissue damage following experimental MCAO (Bederson et al., 1986d). In the present study, SMEF-treated rats exhibited lower neurological scores and traveled a greater total distance than untreated MCAO rats at 24 h after reperfusion (Figs. 3 and 5). Together with the smaller infarct volumes, these findings support an acute protective effect of SMEF. However, total distance traveled reflects overall locomotor activity within the open-field arena and cannot distinguish improved motor capacity from changes in exploratory behavior or general postoperative condition (Seibenhener and Wooten, 2015). Furthermore, the available comparison does not establish whether locomotor activity returned to pre-ischemic or sham levels. These assessments support improved acute functional outcomes but do not establish complete recovery or identify the cellular mechanisms underlying the observed benefit.

SMEF treatment significantly reduced TTC-defined cerebral infarction at 24 h after reperfusion, supporting an acute tissue-protective effect following transient MCAO (Fig. 4). TTC is widely used to identify experimental cerebral infarction with viable tissue stains red, whereas the infarcted zone appear pale or unstained (Bederson et al., 1986b; Liu et al., 2009). Reductions were observed in both edema-corrected infarct volume and infarct percentage. Edema correction reduces the influence of postischemic swelling on infarct estimation (Lin et al., 1993; Swanson et al., 1990), although these two infarct measurements represent related expressions of the same lesion burden rather than independent outcomes. In contrast, the reduction in hemispheric swelling did not reach statistical significance. Thus, the present data support reduced acute infarction but not establish an effect on hemispheric swelling.

Histopathological evaluation provided additional evidence of tissue protection in both the cortex and striatum. Experimental MCAO produces evolving neuronal abnormalities in these regions, including neuronal shrinkage, nuclear pyknosis, cytoplasmic eosinophilia and vacuolation (Rewell et al., 2017). SMEF-treated rats exhibited lower cortical and striatal injury scores than untreated MCAO rats, consistent with the reduction in TTC-defined infarction. Residual abnormalities nevertheless remained after treatment, indicating that tissue injury was attenuated rather than completely prevented. These findings support protection across both regions, although they do not establish whether the striatal benefit resulted from direct electrical effects or secondary consequences of reduced ischemic injury.

Na⁺/K⁺-ATPase is a biologically relevant therapeutic target in cerebral ischemia-reperfusion injury. Huang et al. (2015) reported reduced pump activity in the ischemic penumbra following transient MCAO in rats. Li et al. (2021) further implicated the interaction between Na⁺/K⁺-ATPase and glutamate transporter-1 in protection against cerebral ischemia–reperfusion injury. In the present study, Na⁺/K⁺-ATPase α1 protein abundance was higher in the untreated MCAO rats than in sham-operated rats, and this elevation was attenuated by SMEF treatment. Kim et al. (2013) also reported a transient increase in α1 protein after 2 h of chemical ischemia in primary cultured rat astrocytes, followed by a decrease with prolonged exposure (Kim et al., 2013). Similarly, Kwon et al. (2003) reported enhanced glial Na⁺/K⁺-ATPase α1 immunoreactivity in the gerbil hippocampus following transient forebrain ischemia, with the highest intensity at 4 days, and proposed that this response may contribute to maintaining ionic homeostasis (Kwon et al., 2003). Although this model differs from ours in experimental model and sampling time, their findings provide precedents for ischemia-associated increases in Na⁺/K⁺-ATPase α1 protein abundance or immunoreactivity. Li et al. (2021) reported reduced hippocampal Na⁺/K⁺-ATPase α1 protein abundance at 6 and 24 h after reperfusion following 2 h of MCAO in rats (Li et al., 2021). In contrast, we observed increased α1 abundance in the sampled cortical and striatal tissue at 24 h. differences in the brain regions analyzed may contribute to these divergent findings, but this has not been demonstrated.

GFAP protein abundance was reduced following MCAO and increased with SMEF treatment. Interpretation of this finding requires consideration of the anatomical region and severity of injury. In a permanent MCAO model, Liu et al. (1999) reported loss of GFAP staining within the infarct alongside increased GFAP mRNA expression in peri-infarct regions, demonstrating that the astrocytic response can differ across the ischemic territory. Accordingly, the increase in GFAP associated with SMEF treatment could reflect preservation of GFAP-containing astrocytic structures, altered expression in surviving astrocytes, or difference in the tissue composition of the sampled region. Region-specific immunostaining combined with additional astrocytic and cell-death markers would help clarify this finding.

SMEF treatment also reduced the MCAO-associated increase in NF-κB p65 protein abundance. Previous experimental work has linked changes in NF-κB p65 expression and nuclear localization with inflammatory responses following cerebral ischemia-reperfusion (Li et al., 2023). The present finding is consistent with an effect on inflammation related protein, but it does not directly establish inhibition of NF-κB signaling. The reduced p65 abundance could be a consequence of reduced tissue injury rather than a primary mechanism or protection. Assessment of nuclear and cytoplasmic p65, pathway activation markers would help distinguish these possibilities. We speculate that SMEF may preserve Na⁺/K⁺-ATPase function during ischemia, thereby supporting ionic homeostasis and potentially influencing NF-κB-mediated inflammatory signaling and astrocytic responses. However, whether the observed changes in NF-κB p65 and GFAP are mechanistically linked to pump function remains to be established.

Several limitations should be considered, first, outcomes were assessed only at 24 h after reperfusion. Histopathological changes and behavioral deficits continue to evolve after transient MCAO, supporting the need for longer-term assessment (Rewell et al., 2017). Second, the study included only young adult male rats, limiting generalization to females, aged animals and animals with comorbidities. Further studies are also needed to determine the underlying molecular mechanisms and efficacy of delayed treatment.

Overall, SMEF treatment during the transient MCAO attenuated acute cerebral injury and improved functional outcomes at 24 h after reperfusion. The findings of the present study support the potential of SMEF as a therapeutic approach for ischemic stroke.

## Declarations Ethics

All animal procedures were approved by the Institutional Animal Care and Use Committee of the University of South Florida (protocol no. 11894R) and were conducted in accordance with the National Institutes of Health guidelines for the care and use of laboratory animals.

## Data availability

The data supporting the findings of this study are available from the corresponding author upon reasonable request.

## Supporting information

Supplementary Table S1. Histopathological injury scoring criteria

## Author contributions

R.L. and D.R.C. conceived and designed the study. L.W. and K.B.A. contributed to the experimental design and interpretation of the findings. W.C. and R.L. developed the SMEF platform and stimulation protocol.

D.R.C. implanted the cortical electrodes, performed MCAO surgery, and administered SMEF stimulation.

D.R.C. and N.H. conducted behavioral assessments and quantified infarct volume from TTC-stained brain sections. N.H. and K.W. scored histopathological injury in the cortex and striatum. N.N.Z., Z.T., G.A., and

Y.T. performed Western blotting and densitometric analysis. D.R.C. analyzed the data and prepared the figures. D.R.C. and N.N.Z. drafted the manuscript. R.L. supervised the study. All authors reviewed and approved the final manuscript.

