## Supplementary Table S1. Histopathological injury scoring criteria for "Modified Synchronization Modulation Electric Field attenuates acute brain injury and improves neurological outcomes following stroke induced by transient middle cerebral artery occlusion in rats"

**Supplementary Table S1**. Study-specific semiquantitative scoring criteria for cerebral cortical and striatal histological injury

| Component | Score | Typical features |
| --- | --- | --- |
| Neuronal injury (0-4) | 0 | Normal neurons; no pyknosis; no eosinophilic 'red' neurons |
| | 1 | Occasional pyknotic/red neurons; minimal dropout |
| | 2 | Scattered red neurons + mild neuronal loss |
| | 3 | Many red neurons + moderate loss/necrosis |
| | 4 | Severe necrosis/infarct core |
| Edema/ Vacuolation (0-3) | 0 | None |
| | 1 | Mild vacuolation |
| | 2 | Moderate vacuolation |
| | 3 | Severe diffuse vacuolation |
